# Strand-specific cDNA library-based RNA sequencing dataset of un-infected and *Ascosphaera apis*-infected larval guts of *Apis cerana cerana*

**DOI:** 10.1101/2020.04.09.033738

**Authors:** Huazhi Chen, Zhiwei Zhu, Jie Wang, Yuanchan Fan, Haibin Jiang, Yanzhen Zheng, Cuiling Xiong, Dafu Chen, Rui Guo

## Abstract

*Apis cerana cerana* is a subspecies of eastern honeybee, *Apis cerana*. *Ascosphaera apis* is a widespread fungal pathogen of honeybee, leading to chalkbrood, which results in heavy losses for beekeeping industry. In this article, 4-, 5-, and 6-day-old larval guts of un-infected (AcCK1, AcCK2, and AcCK3) and *A. apis*-infected *A. c. cerana* (AcT1, AcT2, and AcT3) were sequenced by next generation sequencing. Totally, 73830148, 96586212, 94552744, 76672564, 90954858, and 83418832 raw reads were respectively produced from AcCK1, AcCK2, AcCK3, AcT1, AcT2, and AcT3. The sequencing depth was enough to detect all expressed genes. After strict quality control, 73775592, 96513798, 94495000, 76593924, 90870608 and 83339288 clean reads were obtained, with a mean GC content of 48.54%. Additionally, average Q20 and Q30 for aforementioned six groups were 98.10% and 94.36%, respectively. Moreover, 45302685, 65872823, 52709987, 49947838, 56476339, and 42657156 clean reads from above-mentioned six groups were mapped to the reference genome of *Apis cerana*, respectively. In addition, exons were the most abundant regions in reference genome mapped by clean reads, followed by intergenic regions and introns. Our data presented here can be used to identify long non-coding RNAs (lncRNAs), circlular RNAs (circRNAs), mRNAs and their regulatory networks engaged in response of eastern honeybee larvae to *A. apis* infection, and decipher molecular mechanisms underlying host-pathogen interaction.

**Value of the Data:**

- The current data can be used for exploration of mRNAs, lncRNAs, circRNAs and their competing endogenous RNA (ceRNA) networks engaged in response of *A. c. cerana* larvae to *A. apis* infection.
- This data provides a valuable genetic resource for better understanding non-coding RNA-mediated cross-kingdom regulation of eastern honey larvae to *A. apis*.
- The accessible data offers novel insights into understanding the molecular mechanism underlying interaction between eastern honeybee larvae and *A. apis*.

## Data

The shared datasets were derived from strand-specific cDNA library-based RNA sequencing of un-infected (AcCK1, AcCK2, and AcCK3) and *A. apis*-infected (AcT1, AcT2 and AcT3) *A. c. cerana* 4-, 5-, and 6-day-old larval guts. In total, 73830148, 96586212, 94552744, 76672564, 90954858, and 83418832 raw reads were gained from AcCK1, AcCK2, AcCK3, AcT1, AcT2, and AcT3, respectively (**Table 1**). Additionally, the sequencing depth was enough to detect all expressed genes (**Figure 1**). After strict quality control, 73775592, 96513798, 94495000, 76593924, 90870608 and 83339288 clean reads were respectively gained from the six groups mentioned above (**Table 1**). Besides, average Q20, Q30 and GC content were 98.10%, 94.36% and 48.54%(**Table 2**). Moreover, 45302685, 65872823, 52709987, 49947838, 56476339, and 42657156 clean reads were mapped to the reference genome of *Apis cerana*, respectively (**Table 3**). As shown in Table 4, exons were the most abundant regions in reference genome mapped by clean reads, followed by intergenic regions and introns.

**Table 1.**
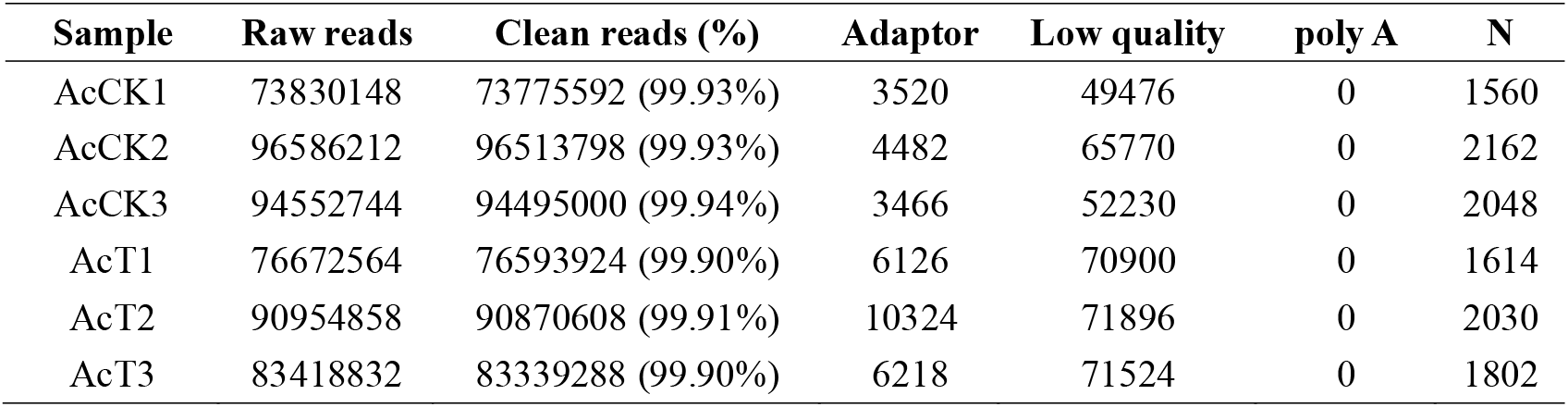
A summary of filtering of raw reads from strand-specific cDNA library-based RNA sequencing.

| **Sample** | **Raw reads** | **Clean reads (%)** | **Adaptor** | **Low quality** | **poly A** | **N** |
| --- | --- | --- | --- | --- | --- | --- |
| AcCK1 | 73830148 | 73775592 (99.93%) | 3520 | 49476 | 0 | 1560 |
| AcCK2 | 96586212 | 96513798 (99.93%) | 4482 | 65770 | 0 | 2162 |
| AcCK3 | 94552744 | 94495000 (99.94%) | 3466 | 52230 | 0 | 2048 |
| AcT1 | 76672564 | 76593924 (99.90%) | 6126 | 70900 | 0 | 1614 |
| AcT2 | 90954858 | 90870608 (99.91%) | 10324 | 71896 | 0 | 2030 |
| AcT3 | 83418832 | 83339288 (99.90%) | 6218 | 71524 | 0 | 1802 |

**Table 2.**
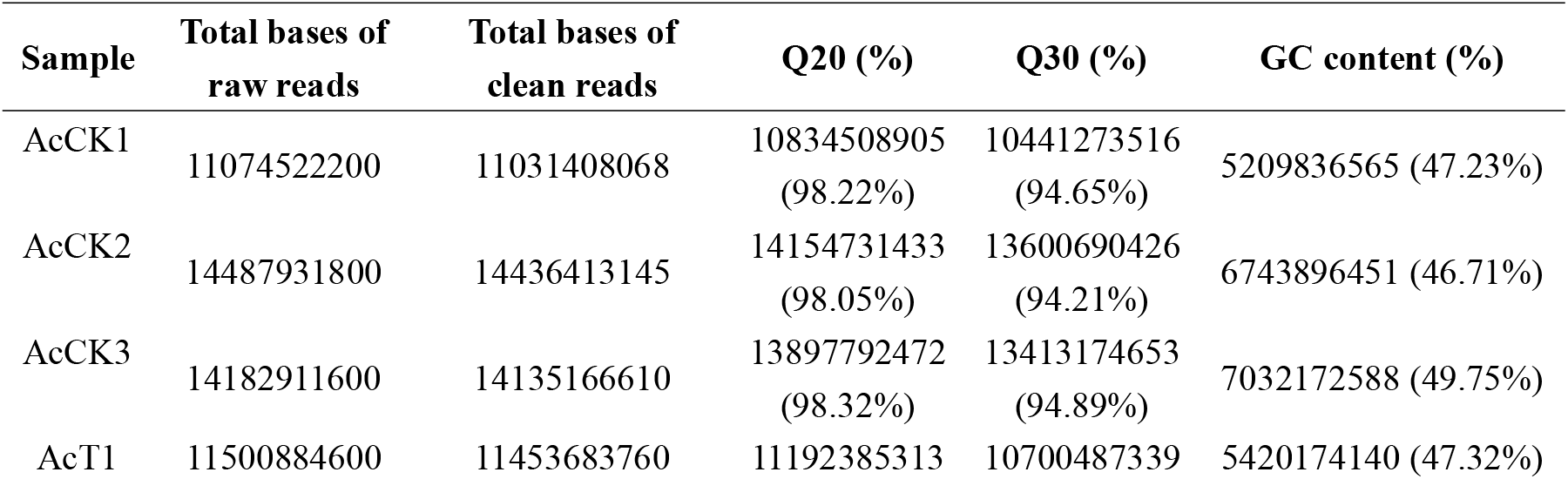
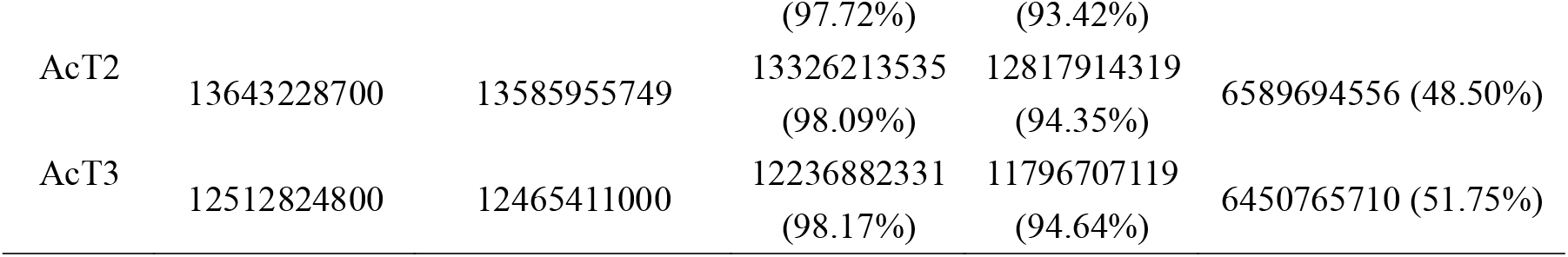
An overview of clean reads after quality control.

**Table 3.**
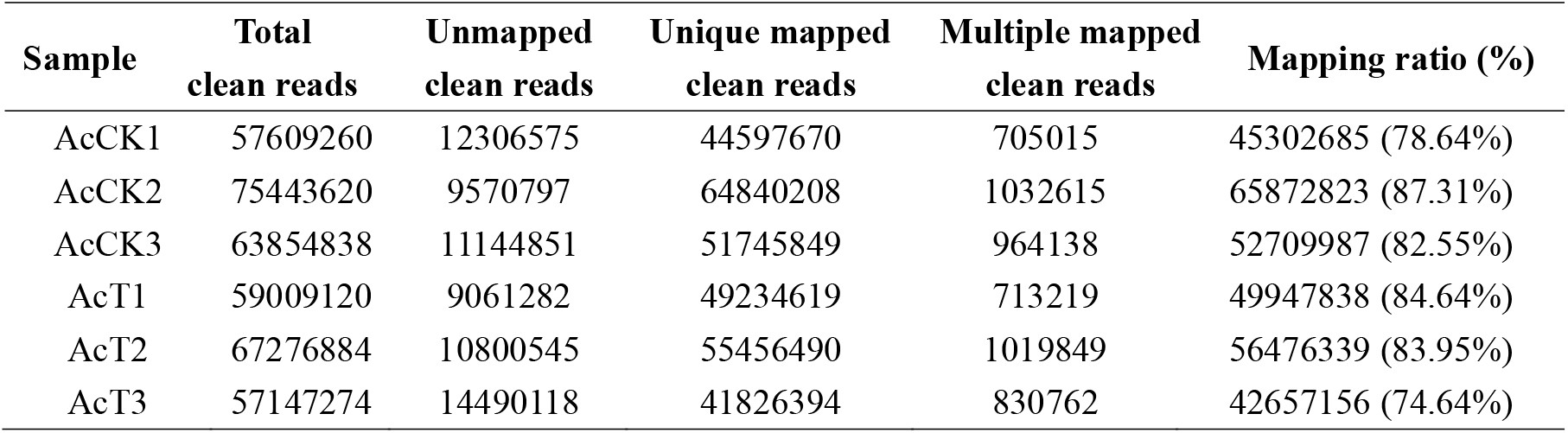
A summary of mapping of clean reads to reference genome of *A. cerana*.

| Sample | Total<br>clean reads | Unmapped<br>clean reads | Unique mapped<br>clean reads | Multiple mapped<br>clean reads | Mapping ratio (%) |
| --- | --- | --- | --- | --- | --- |
| AcCK1 | 57609260 | 12306575 | 44597670 | 705015 | 45302685 (78.64%) |
| AcCK2 | 75443620 | 9570797 | 64840208 | 1032615 | 65872823 (87.31%) |
| AcCK3 | 63854838 | 11144851 | 51745849 | 964138 | 52709987 (82.55%) |
| AcT1 | 59009120 | 9061282 | 49234619 | 713219 | 49947838 (84.64%) |
| AcT2 | 67276884 | 10800545 | 55456490 | 1019849 | 56476339 (83.95%) |
| AcT3 | 57147274 | 14490118 | 41826394 | 830762 | 42657156 (74.64%) |

**Table 4.** An overview of mapped region of clean reads in *A. cerana* genome.

| Sample | Exon (%) | Intron (%) | Intergenic region (%) |
| --- | --- | --- | --- |
| AcCK1 | 30417508 (67.14%) | 5319947 (11.74%) | 9565230 (21.11%) |
| AcCK2 | 43199223 (65.58%) | 5896658 (8.95%) | 16776942 (25.47%) |
| AcCK3 | 34629762 (65.70%) | 3934167 (7.46%) | 14146058 (26.84%) |
| AcT1 | 31818489 (63.70%) | 5101262 (10.21%) | 13028087 (26.08%) |
| AcT2 | 37231677 (65.92%) | 4795495 (8.49%) | 14449167 (25.58%) |
| AcT3 | 26578893 (62.31%) | 3306117 (7.75%) | 12772146 (29.94%) |

**Figure 1.**
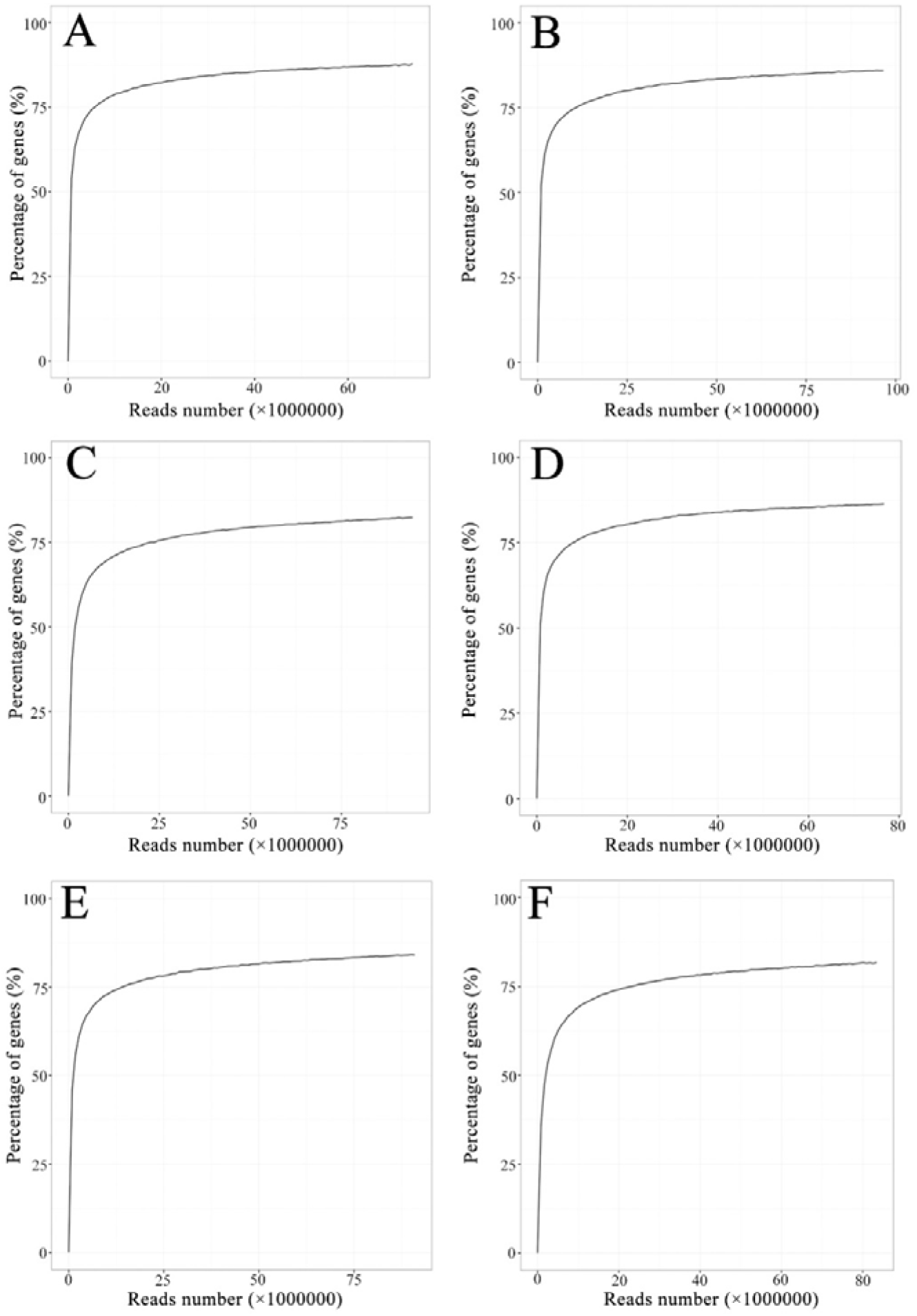
Sequencing satuation of six sample groups. A: AcCK1 group; B: AcCK2 group; C: AcCK3 group; D: AcT1 group; E: AcT2 group; F: AcT3 group.

## Experimental Design, Materials, and Methods

### 2.1 Fungal spore purification

*A. apis* [1–2] was previously isolated from a fresh chalkbrood mummy according to the method described by Jensen et al [3], and kept in Honeybee Protection Lab in College of Animal Sciences (College of bee Science), Fujian Agriculture and Forestry University. Following the protocol developed by Jensen [3] with some minor modifications [4], fresh spores of *A. apis* were purified at 7 days after culturing under lab condition.

### 2.2 Inoculation of honeybee larvae and preparation of gut sample

*A. c. cerana* larvae were artificially reared according to the previous method [5]. The artificial diet was mixed and frozen in smaller aliquots, and pre-heated to 34 ^°^C before use. Two-day-old (post-hatch) larvae were carefully removed from the combs using a Chinese grafting tool and transferred to a droplet of 10 μL diet in each well in sterile 48-well culture plates. Larvae were fed once a day with 20 μL (3-day-old), 30 μL (4-day-old), 40 μL (5-day-old) and 50 μL (6-day-old) diet.

To cause effective infection, 3-day-old larvae in treatment groups were fed with diet containing *A. apis* spores at a final concentration of 10^7^ spores/mL [6]. Three-day-old larvae in control groups were fed with diet without fungal spores. Hereafter, larvae in control and treatment groups were reared with normal diet. Culture plates were incubated at 95% RH and 33 °C [7]. Following our previously developed method [6], larval guts of 4-, 5- and 6-day-old larvae (n=7) were harvested from *A. apis*-infected and un-infected groups. These larval gut samples were frozen immediately in liquid nitrogen and stored at −80 °C until next-generation sequencing. Gut samples selected from 4-, 5-, and 6-day-old larvae in *A.apis*-infected groups were termed as AcT1, AcT2, and AcT3; while those selected from 4-, 5-, and 6-day-old larvae in un-infected groups were termed as AcCK1, AcCK2, and AcCK3.

### 2.3 Strand-specific cDNA library construction and next-generation sequencing

Following the manufacturer’s protocol, total RNA of the larval guts from each *A. apis*-infected group and un-infected group were respectively isolated using Trizol (Life Technologies), and then examined via 1% agarose gel eletrophoresis. Next, mRNAs and ncRNAs were retained after removal of rRNAs, and then fragmented into short fragments with fragmentation buffer (Illumina, USA) followed by reverse transcription into cDNAs with random primers. Second-strand cDNAs were synthesized by dNTP (dUTP instead of dTTP), DNA polymerase I, RNase H, and buffer. By using QiaQuick PCR extraction kit (QIAGEN, Germany), the cDNA fragments were purified, end repaired, poly(A) added, and ligated to Illumina sequencing adapters, followed by digestion of the second-strand cDNAs with UNG (Uracil-N-Glycosylase) (Illumina, USA). Ultimately, the digested products were size selected via agarose gel electrophoresis, PCR amplified, and sequenced on Illumina HiSeq™ 4000 platform (Illumina, USA) by Gene Denovo Biotechnology Co. (Guangzhou, China).

### 2.4 Quality control of raw reads and mapping of clean reads

To obtain high quality clean reads, raw reads yielded from Illumina sequencing were first filtered by removing reads that contain adapters, more than 10% of unknown nucleotides (N), and more than 50% of low quality bases. Information about quality control of raw data is summarized in **Table 1**. Next, clean reads were mapped to ribosome RNA (rRNA) database (http://rdp.cme.msu.edu/) using short reads alignment tool Bowtie2 [8]. The mapped clean reads were removed and the remaining clean reads were used for transcript assembly and further analysis. Subsequently, the rRNA-removed clean reads of each sample group were mapped to the reference genome of *Apis cerana* (assembly ACSNU-2.0) with TopHat2 (version 2.0.3.12) [9] following alignment parameters: maximum read mismatch is two; the distance between mate-pair reads is 50 bp; the error of distance between mate-pair reads is ± 80 bp.

## Acknowledgments

This research was financially supported by the National Natural Science Foundation of China (No. 31702190), the Earmarked Fund for China Agriculture Research System (No. CARS-44-KXJ7), the Science and Technology Planning Project of Fujian Province (No. 2018J05042), the Teaching and Scientific Research Fund of Education Department of Fujian Province (No. JAT170158).

